# Drug-eluting biodegradable metals and metal-ceramic composites: High strength and delayed drug release

**DOI:** 10.1101/2022.12.22.521630

**Authors:** Aliya Sharipova, Olga Bakina, Aleksandr Lozhkomoev, Marat Lerner, Elazar Gutmanas, Alejandro Sosnik

## Abstract

Biodegradable metals emerged as promising temporary bone implants. The integration of additional features such as local drug delivery (LDD) can also support their osteointegration, promote bone regeneration, and prevent biomaterial-centered infections that are difficult to treat. LDD is achieved by drug-eluting coatings or porous implants where the drug is impregnated after implant fabrication because the high temperatures used during conventional production processes would result in their thermal decomposition. We produced biodegradable iron (Fe)-based vancomycin (VH)-eluting metals and metal-ceramic composites by a simple high-pressure consolidation/cold sintering (CS) process at room temperature that display high mechanical strength and antibacterial activity. Aiming to expand the application of this production method and shed light into the drug loading and release mechanisms in this type of biomaterials, this work reports on the production and characterization of VH-loaded Fe and Fe-iron oxide (Fe_2_O_3_) composites (Fe-Fe_2_O_3_). We use focus ion beam milling for the first time to investigate the drug-metal interface and investigate the mechanical and degradation properties of VH-free and VH-loaded Fe and Fe-Fe_2_O_3_. Results show very high mechanical strength of drug-eluting Fe and Fe-Fe_2_O_3_ composites (up to than 780 MPa under compression, exceeding the maximum strength of cancellous bone more than three times) accompanied by a delayed drug release. Then, we confirm the good antimicrobial activity against *Staphylococcus aureus* and cell compatibility with the murine embryonic fibroblast cell line NIH/3T3 *in vitro*. Overall results confirm the promise of drug-eluting metals and metal-ceramic composites for LDD in bone.

## 1. Introduction

Biodegradable orthopedic implants are an attractive strategy in the healing of critical-size bone defects because they provide temporary mechanical support at the healing site leveraging the ability of this tissue for self-restoration and are gradually replaced by the regenerating tissue [1,2]. Biodegradable implants also preclude the need for an explantation surgery improving patient compliance [3], shortening the recovery time, reducing operational costs, labor load, and intervention-related risks [4,5].

To date, a broad spectrum of biodegradable orthopedic biomaterials has been developed [6]. However, not all of them combine biodegradability with high mechanical strength, ductility and structural stability that are key features for applications in load-bearing bones (e.g., pelvis, tibia, femur, knee joint) [7]. Among the existing biomaterials, only biodegradable metals or metal-based composites (e.g., metal-metal and metal-ceramic) can fulfill these mechanical requirements [8,9] as biodegradable polymers do not provide adequate mechanical support at load-bearing sites, while biodegradable ceramics are naturally brittle. In this scenario, biodegradable metals are promising candidates for temporary load-bearing bone implants [9,10].

Bone- and implant-related infections such as osteomyelitis remain a substantial burden in healthcare showing increasing trend over the last decades especially among the old population. According to comprehensive population-based study in the 1969-2009 period, the annual incidence of osteomyelitis almost tripled among American individuals older than 60 years [11]. Among the cases, 44% were caused by *Staphylococcus aureus* (*S. aureus*) and 17% *by Staphylococcus epidermidis* [11]. Cross-sectional study of vertebral osteomyelitis epidemiology reported an increase of ~59% from 2010 to 2019 among French adults (mean age of 64.8 years old) [12]. Similar to the US studies, a major source of vertebral osteomyelitis among the French individuals was the *Staphylococci* family resulting in 35.2% of all cases [12]. Bone-related infections are especially difficult to treat via systemic antibiotic administration due to more limited blood supply in bone than in soft tissues [13]. In this conceptual framework, the integration of local drug delivery (LDD), namely the release of active compounds at the body site of interest, features that prevent infections and/or promote osteointegration and bone regeneration can reduce the risk of post-operative complications and increase the success rates. LDD is more advantageous than systemic drug administration because it reduces the effective drug dose, ensures maximum bioavailability in a tissue with more limited blood supply and minimizes associated systemic side-effects [14–16].

LDD features can be achieved in biodegradable metallic implants by the integration of porous coatings to bulk biomaterials [17] or by the production of porous implants/scaffolds where the drug is infiltrated or impregnated post-production by different methods [18,19]. Drug-eluting coatings typically have an initial burst release followed by a slower release rate for a few days owing to their limited loading capacity [14,20]. Such release kinetics is usually sufficient to prevent or treat intervention-related infections. However, in some applications, extended or delayed release of bioactive substances such as bone morphogenetic proteins (BMPs) that result in a more efficient and stable osteointegration at the bone-implant interface might be preferred [21–23]. Drug-infiltrated/impregnated porous metals prolong release kinetics because of a greater loading capacity though they display lower mechanical strength than bulk counterparts [18], jeopardizing their use in LDD at load-bearing bones. To ensure adequate mechanical support at the healing site during implant degradation, extended LDD requires drug loading into the implant bulk. Conversely, the fabrication of drug-loaded bulk metals is challenging because of the high temperatures used in metal processing that leads to thermal drug decomposition. Recently, we reported on the production of biodegradable vancomycin (VH)-eluting iron-silver (Fe10Ag) bulk metals by a simple high-pressure consolidation/cold sintering (CS) process at room temperature (RT) that display good mechanical strength and antibacterial activity [24]. These biodegradable drug-eluting metals showed mechanical strength comparable to that of cortical bone and extended drug delivery during five days [24]. At the same time, their yield strength under compression (180 MPa) and bending (118 MPa) was 4- and 8-fold lower, respectively, than drug-free Fe10Ag counterparts [25]. Although, this detrimental effect of the loaded drug in the mechanical properties of the metal was anticipated, further studies are called for to characterize the drug-metal interface and understand the fundamental parameters that govern the mechanical performance of drug-eluting metals. Based on our previous experience with biodegradable metals produced by CS [25–28], we hypothesized that the fine tuning of the microstructure of the metallic matrix would lead to better consolidation during CS, and to improved mechanical features.

Aiming to shed light into the microstructure, and drug loading and release mechanisms in CS-produced drug-eluting metals and to expand the application of this production method, in this work, we fabricate and comprehensively characterize VH-free and VH-loaded metals (Fe) and metal-ceramic composites (Fe with iron oxide, Fe-Fe_2_O_3_) with a mechanical strength that exceeds by more than three times the maximum strength of cancellous bone (~220 MPa in longitudinal direction) [29,30]. Overall results show the potential of CS to produce mechanically strong drug-eluting metal and metal-ceramic implants.

## 2. Materials and Methods

### 2.1. Preparation of VH-free and VH-loaded Fe and Fe-Fe_2_O_3_

Fe-Fe_2_O_3_ (5, 10, 15 vol% of Fe_2_O_3_) powder blends were prepared by manual mixing of carbonyl Fe powder and Fe_2_O_3_ nanopowder. Carbonyl Fe powder (5-9 μm) was purchased from STREM Chemicals (Newburyport, MA, USA). Fe_2_O_3_ nanopowders (50-200 nm) was chemically synthesized as reported elsewhere [31]. For preparation of Fe-5Fe_2_O_3_ blends, 0.5872 g of Fe and 0.0206 g of Fe_2_O_3_ were weighed in a precision digital balance ES120A (Precisa, Dietikon, Switzerland). To ensure the homogeneous mixing of the powders, each powder was roughly divided into five fractions, and one fraction of each component were dry-mixed in an agate mortar with pestle. The same procedure was repeated for all the fractions. Afterwards, all the Fe-Fe_2_O_3_ mixtures were combined and dry-mixed together. This mixing procedure was employed for preparing other Fe-Fe_2_O_3_ blends (with 10 and 15 vol% of Fe_2_O_3_).

In this work, VH (STREM Chemicals) was used as a model antibiotic drug due to its wide use for the treatment of periprosthetic and implant-related infections [32]. To prepare Fe samples loaded with 1 wt% of VH (Fe-1wt%VH), 0.6000 g of the Fe was mixed with 0.0061 g of VH powder following the mixing procedure described above. Then, the Fe with 1wt% VH precursor mixture was loaded into a custom-build consolidation die and high-pressure cold-sintered at 2.5 GPa (corresponding to 5 t for 5 mm diameter die) and RT using manual press (Carver, Wabash, IN, USA) to obtain the drug-loaded metal (Supplementary Materials, **Fig. S1**). A similar procedure was utilized to prepare Fe-5Fe_2_O_3_ and Fe-10Fe_2_O_3_ samples loaded 1 wt% VH (Fe-5Fe_2_O_3_-1wt%VH and Fe-10Fe_2_O_3_-1wt%VH, respectively), and a representative sample of Fe containing 5 wt% VH for the microstructure investigations.

### 2.2. Structural and mechanical characterization of VH-free and VH-loaded Fe and Fe-Fe_2_O_3_

The microstructure of the different materials was characterized by means of high-resolution scanning electron microscopy (HR-SEM, Ultra+, Zeiss, Oberkochen, Germany) equipped with back-scatter electron (BSE) detector. Cross-sectional micrographs were obtained by focused ion beam scanning electron microscope (FIB, Helios NanoLab, FEI, Thermo Scientific, Waltham, MA, USA) using Everhart-Thornley secondary electron detector (ETD) or through-lens (Elstar in-lens) secondary electron detector (TLD) and energy-dispersive spectroscopy (EDS) detector. Platinum coating (Pt coating) was deposited to protect the zone of interest during destructive FIB milling. FIB slicing technique was employed to prepare three-dimensional micrograph (3D-micrograph). A volume 25 × 25 × 25 μm of Fe with 5 wt% VH sample was cut into slices with 100-nm steps using Auto Slice&View 1.7 Software (Thermo Scientific). A 3D-micrograph was reconstructed based on the set of collected images (near 250 HR-SEM cross-sectional micrographs) in Avizo 9.5 Software using Vortex module (Thermo Scientific).

The sample density was measured by the Archimedes method and the Relative density (*Rel. ρ*, expressed as %) was calculated according to **Equation 1**

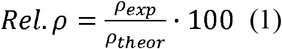

where *ρ_exp_* is sample density (g/cm^3^) measured by Archimedes method and *ρ_theor_* is the sample theoretical density. We considered VH density (1.65 g/cm^3^) and content in calculations of theoretical density of drug-eluting samples.

The mechanical properties of the different materials were characterized under compression using a universal testing machine (Instron 5900, Instron, Norwood, MA, USA) at a strain rate of 200 μm/min.

### 2.3. Degradation of Fe and Fe-Fe_2_O_3_

Static degradation tests of VH-free Fe and Fe-Fe_2_O_3_ (5, 10, 15 vol% Fe_2_O_3_) samples were conducted under immersion according to the ASTM G31 standard [33]. Samples (5 mm in diameter × 4 mm in height) were immersed in modified Hanks’ solution (MH) prepared as described elsewhere [34]. Briefly, 9.7□g of Hanks’ balanced salt (Sigma-Aldrich, St. Louis, MO, USA) was dissolved in 1.4□L of MilliQ water (Barnstead Smart2Pure 3 L UV/UF water purification system, Thermo Scientific) with 14.16□g of HEPES acid (STREM Chemicals) and 16.65 g of HEPES sodium salt (STREM Chemicals). The solution pH was adjusted to pH 7.4 ± 0.1 with 1□M sodium hydroxide (NaOH, Bio-Lab Ltd., Jerusalem, Israel) and/or 1□M hydrochloric acid (HCl, Bio-Lab Ltd.). Each sample was suspended separately by a nylon mesh in the middle of a 50-mL conical polypropylene Falcon tube with 50 mL of MH and a magnetic stirrer. This MH volume satisfies the minimum required by ASTM G31 standard (0.40□mL/mm^2^ “solution volume-to-specimen area” ratio) [33]. Samples were incubated in a water bath at 37□±□1 °C under constant magnetic stirring for eight weeks. Every seven days, samples were extracted, immersed three times in ultrasound bath of fresh 70% ethanol solution for 10□min to remove corrosion products, dried in a desiccator under vacuum overnight and gently cleaned with a soft brush to remove remaining corrosion products. The weight of the sample and pH of the collected aliquots were measured. To continue the degradation test, samples were placed in fresh pre-heated 50 mL of MH as was described above and incubated until the next aliquot extraction.

The degradation rate (expressed in mm/year) was calculated from the weight loss measurements, according to **Equation 2**

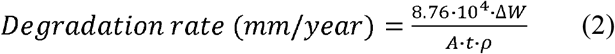

where Δ*W* is weight change (g), *A* is the surface area (cm^2^), *t* is the immersion time (h) and *ρ* is the initial density of the sample (g/cm^3^).

### 2.4. Antibacterial activity of VH-free and VH-loaded Fe and Fe-10Fe_2_O_3_

The antimicrobial activity of VH-loaded (1 wt%) Fe and Fe-10Fe_2_O_3_ composites was assessed by the zone of inhibition (ZOI) test, according to the procedure described elsewhere [35]. In brief, each sample was immersed in sterile sodium chloride solution (1 mL, 0.9% w/v, Grotex, St. Petersburg, Russia) in 5-mL Falcon tubes (Corning, Glendale, AZ, USA) and incubated at 37 °C and 95% humidity. At different time points, aliquots (100 μL) were taken, replaced with sodium chloride solution and stored at −20 °C until analysis. Then, the antibacterial activity against *S. aureus* strain ATCC 6538 (BRC VKM, Moscow, Russia) was tested. Bacteria were grown in Miller-Hinton broth (Sigma-Aldrich), suspensions with a McFarland standard of 0.5 were spread onto Mueller-Hinton agar plates (Heipha Dr. Müller GmbH, Eppelheim, Germany), test filter discs (Ø 6 mm, HiMedia Laboratories Pvt Ltd., Mumbai, India) were placed onto the agar and 15 μL of each elution sample was pipetted onto them. Agar plates were incubated overnight at 37 °C and the ZOI (in mm) was measured.

### 2.5. Cell compatibility of Fe and Fe-10Fe_2_O_3_

The mouse embryonic fibroblast cell line NIH/3T3 (ATCC CRL-1658, supplied by the State Research Center of Virology and Biotechnology “VECTOR”, Koltsovo, Russia) was used to evaluate the cell compatibility of Fe and Fe-10Fe_2_O_3_ composites. For this, cells were cultured in Dulbecco’s Modified Eagle Medium (DMEM, HyClone Laboratories, Inc., Logan, UT, USA) with the addition of 10% fetal bovine serum (HyClone) and 1% penicillin/streptomycin gentamicin (HyClone) at 37 °C and 5% CO_2_ within 24 h until cell confluence was reached. Then, cells were washed with Dulbecco’s Phosphate Buffered Saline (DPBS, Sigma-Aldrich) and harvested using 0.25% trypsin-EDTA (Sigma-Aldrich). Cells were counted by a TC20 Automated Cell Counter (Bio-Rad Laboratories, Inc., Hercules, CA, USA).

To rule out the possible toxic effect of VH, only the cell compatibility of drug-free Fe and Fe-10Fe_2_O_3_ specimens was assessed. Cells (70,000 cells per well) were cultured in 24-well plates (total culture medium volume of 2 mL) for 48 h in the presence of Fe and Fe-10Fe_2_O_3_ samples. After incubation, cells were washed with DPBS and harvested with trypsin-EDTA and the viability analyzed by flow cytometry using the Apoptosis Detection Kit with propidium iodide (PI, BioLegend, San Diego, CA, USA) in a Cytoflex flow cytometer (Beckman Coulter, Brea, CA, USA). Results were processed using CytExpert 2.0 software (Beckman Coulter). For statistical significance, at least 10,000 cells were analyzed in each sample [36]. Untreated cells were used as control and considered 100% viable.

### 2.7 Statistical analysis

Statistical analysis of the degradation rate was performed by t-test on raw (relative weight loss) data using “Two-Sample t-test on Rows” tool in OriginPro 2022 software (OriginLab Corporation, Northampton, MA, USA). p-values of less than 0.05 were considered as statistically significant. Mechanical testing results were expressed as Mean ± standard deviation (S.D.) of three samples (n = 3). ZOI results are expressed as the Mean ± S.D. of five replicates (n = 5). For each sample, cell compatibility assay was performed five times (n = 5).

## 3. Results and Discussion

### 3.1. The concept

Previously, we reported on the fabrication of dense drug-eluting metals from nanostructured Fe10Ag powders using CS at RT [24]. However, use of nanostructured powders might be disadvantageous when consolidation to full density is desired [37]. Increased number of grain boundaries can obstruct plastic flow of metallic particles, which is required for the consolidation during CS. The extent to which particles plastically deform depends on the microstructure and follows the Hall-Petch rule [38]

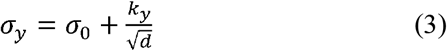

where *σ_y_* is the yield stress of the metal, *σ*_0_ is the resistance of the lattice to dislocation motion, *k_y_* is the strengthening coefficient (a constant specific to each material) and *d* is the average grain diameter of the material. Given that *σ*_0_ and *k_y_* are constant for a specific material, *σ_y_* or the external pressure required to plastically deform the particle depends on its grain size *d*. At the same pressure, microstructured particles (*d* ~ 100 μm) would display a greater plastic deformation/strain, so provide close contact between neighboring particles during CS than their nanostructured counterparts (*d* ~ 100 nm). Since there are no other means of matter transport during the CS (e.g., due to the heat treatment), microstructured particles can provide larger strains and undergo better consolidation under the applied pressure. Therefore, we hypothesize that microstructured powders can improve the densification of drug-eluting metals by CS, so the mechanical performance. For this purpose, we used carbonyl Fe powder (**Fig. 1**), which grains are at the micron- and sub-micron scale (**Fig. 1c**). Shades of grey correspond to orientational (grain) contrast in cross-section of the Fe particle (**Fig. 1b,c**).

**Fig. 1.**
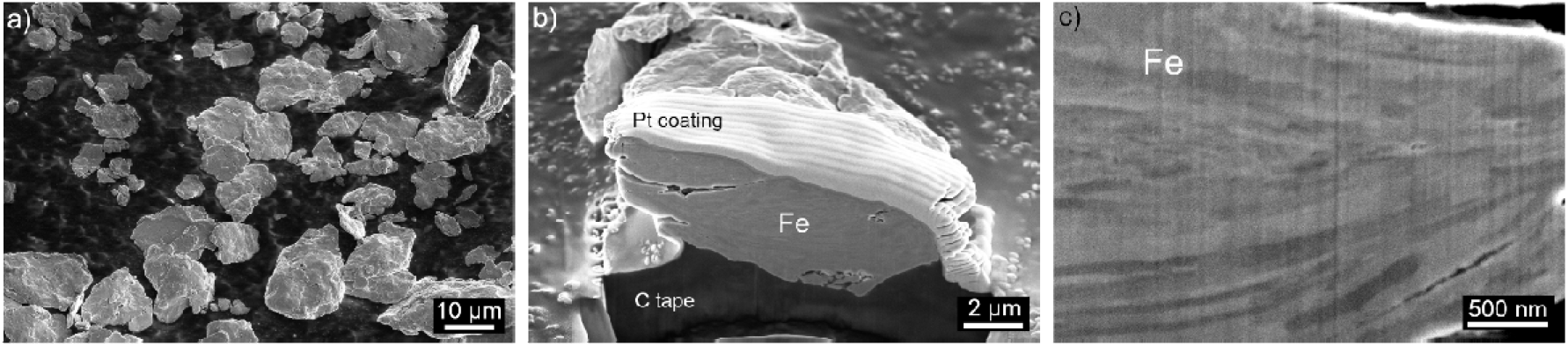
HR-SEM micrographs of carbonyl Fe powder: (a) as-provided; (b,c) cross-sectional images of a representative Fe particle. Pt coating – protective coating for FIB milling. C tape – carbon conductive tape.

Considering the low degradation rate of Fe for biodegradable applications [8], we introduced Fe_2_O_3_ nanopowder (10 ± 5 vol%), which was reported to increase degradation rate of the Fe matrix [39]. In this work, we denote Fe samples as “metal” and Fe-Fe_2_O_3_ samples as “metal-ceramic composites” (also “metal-ceramics” or “composites”). VH-loaded Fe is denoted as “drug-eluting metal”, and VH-loaded Fe-Fe_2_O_3_ as “drug-eluting metal-ceramic composites” (shortly “drug-eluting metal-ceramics”, or “drug-eluting composites”). At the same time, it is important to emphasize that primary phase in VH-loaded Fe-Fe_2_O_3_ is metallic Fe (90 ± 5 vol%), which enables consolidation and drug loading at the given conditions. In this framework, VH-loaded metal and metal-ceramic composites were produced by a CS method that ensures optimal drug encapsulation efficiency, homogeneous distribution without thermal decomposition, as previously reported by our group [24]. To ensure drug homogeneity, a powder mixing process was developed (**Fig. S1**).

### 3.2. VH loading into bulk Fe

One of the fundamental questions that remains unraveled is the mechanism of drug encapsulation in bulk metals. Aiming to shed light into this mechanism that has implications in the biodegradation and drug release, we prepared a Fe sample loaded with 5 wt% of VH (Fe-5wt%VH) that corresponds to approximately 25 vol% based on the metal-drug density ratio. Higher amount of drug provides larger area of drug-matrix interface, which helps in HR-SEM characterization. Fe-5wt%VH was processed by FIB and analyzed by HR-SEM to reconstruct the 3D-microstructure. The isometric view of the 3D-micrograph and representative cross-sectional micrographs of Fe-5wt%VH are presented in **Fig. 2**. Here, the light-grey and dark-grey color correspond to the Fe matrix and the encapsulated VH, respectively. The white phase on the top and at the edge of the reconstructed volume (**Fig. 2a**, **Supplementary Video 1**) corresponds to a Pt protective coating used in FIB milling. Vertical texturing of the cross-sectional micrographs (**Fig. 2b-d**) represents the so-called “curtaining effect” in FIB milling. More details on the effect are provided in **Fig. S2** of Supplementary Materials.

**Fig. 2.**
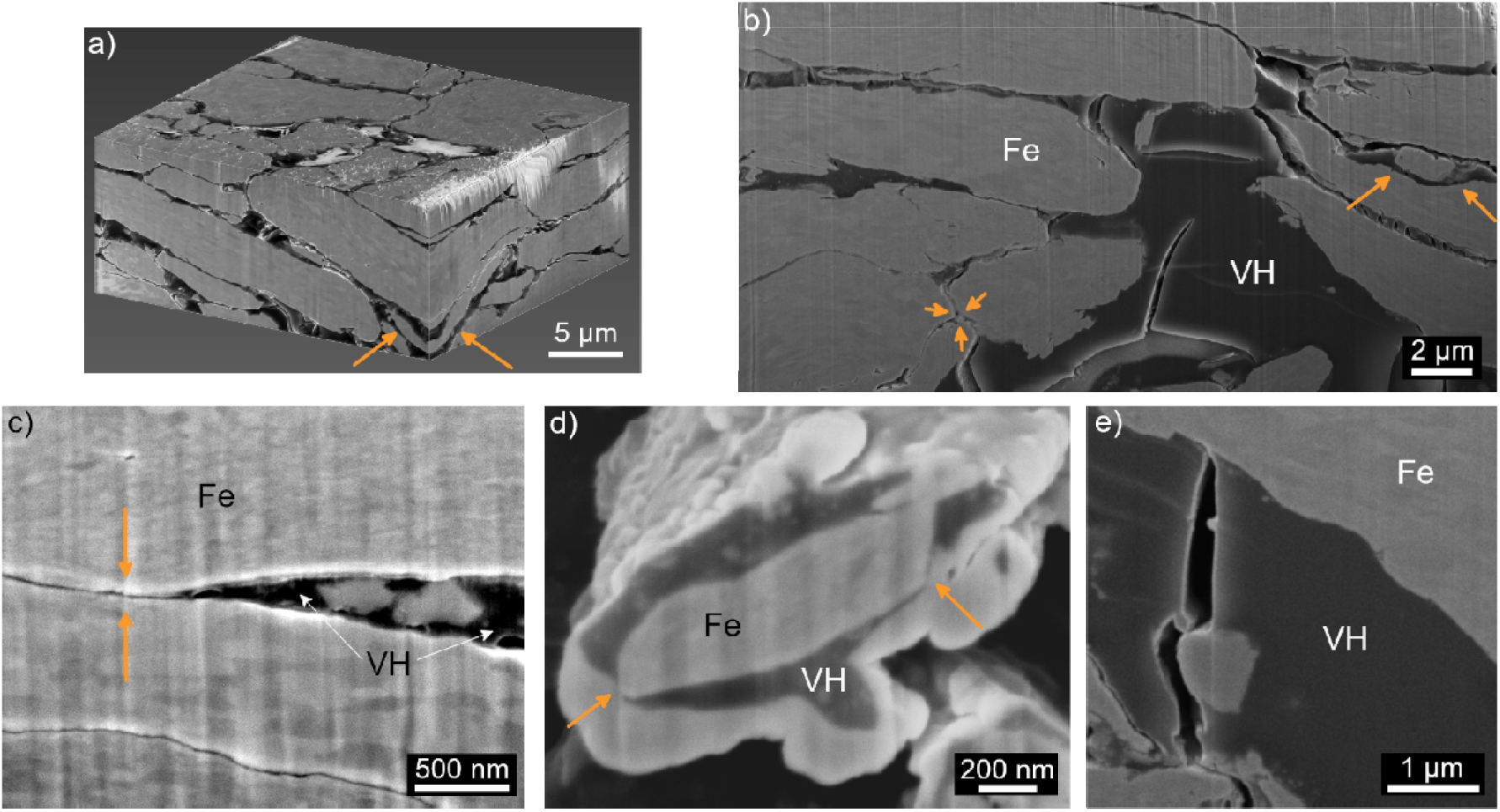
HR-SEM micrographs (TLD) of Fe-5wt%VH produced by the CS method. The encapsulated VH and Fe are visualized as the dark grey and the light grey phases, respectively. (a) 3D-Micrograph: the white phase on the top and the edges is a Pt protective coating used in FIB milling; and (b-e) representative cross-sectional micrographs obtained in different regions of the VH-loaded sample. Yellow arrows in the micrographs point to the signs of (a,b) plastic deformation and (c,d) sintering in Fe particles.

In general, the 3D-reconstruction shows an almost dense metallic matrix enclosing a network of drug particles having a flat sub-micron scale structure (**Fig. 2a**). Such network structure is explained by the flat shape of the initial Fe particles (**Fig. 1a**) and their plastic deformation under pressure. In some regions, the drug network broadens and forms micron-scale inclusions (large black phase in **Fig. 2b**) that are of a similar size as the granules described in our previous work on Fe10Ag loaded with 1 wt% VH [24]. As we observe, an increase of VH content from 1 to 5 wt% (that corresponds to an increase from 5 to 25 vol%) changes its microstructure from a system of separate granules embedded in the metallic matrix [24], to an interconnected network (**Fig. 2a**, **Supplementary Video 1**). This change could be explained by the drug distribution that depends on the content and the drug-metal mixing procedure. In both works, we employed manual mixing in mortar and pestle for the drug loading (**Fig. S1**). Soft drug particles spread over Fe particles by rubbing motion and a greater drug content leads to the formation of a drug network within the metallic matrix upon consolidation. Other drug loading procedures (e.g., dry or liquid mixing, electromagnetic or mechanical stirring [40]) might change the spreading of drug over metallic particles, so the shape and the distribution of the drug in the cold-sintered matrix.

HR-SEM micrographs of the Fe-5wt%VH cross-section show the metallic, drug phases and their interface (**Fig. 2c-e**). Close analysis of the Fe phase reveals no change of the grain structure (**Fig. 2c**), as compared to the initial one (**Fig. 1c**). Cross-sectional micrographs enable tracing the plastic deformation of the metallic phase utilized for the preparation of the drug-eluting metals. First, the surfaces of neighbouring Fe particles replicate each other shapes (yellow arrows in **Fig. 2a,b**). Second, porosity at Fe grain boundaries reflects that they are newly formed between former separate Fe particles (yellow arrows in **Fig. 2c**). Third, the VH phase enclosed between sintered metallic phases is a result of plastically deformed adjacent Fe particles (yellow arrows in **Fig. 2c,d**).

For the first time, we investigated the microstructure of the VH phase (dark phase in **Fig. 2**) and of the Fe-VH interface consolidated at 2.5 GPa and RT. Close examination unveils that, under these conditions, the VH powder forms a dense phase of irregular shape with numerous cracks (**Fig. 2b,e**, **Supplementary Video 1**). The morphology of the cracks is similar to those previously observed upon fracture of Fe10Ag-1wt%VH samples [24]. In this work, the FIB slicing technique enabled to unequivocally establish that cracks are inherent to the VH phase in bulk metal produced by CS. High applied pressures can initiate cracks in the drug phase, which has weak and brittle or semi-brittle nature [41]. Moreover, significant difference in the elastic behavior of Fe and VH contributes to crack development. Young’s modulus of pure Fe ranges between 187 and 214 GPa [42]. While that of VH has not been reported, it can be roughly estimated from the values of different organic pharmaceutical excipients such as starch and microcrystalline cellulose that are <10 GPa [43–45]. Such a ~20-fold difference in the elastic behavior between Fe and VH particles initiates stress concentrations in the mechanically weak drug phase during the CS. Elastic strain of Fe phase later is overtaken by severe plastic deformation. However, elastic strain in Fe matrix is also present after the pressure release during the matrix recovery-relaxation [46,47]. Relaxation of the metallic matrix can initiate or develop existing cracks in the VH phase that we observe in the cold-sintered sample (**Fig. 2b,e**).

Close analysis of HR-SEM micrographs shows no pores at the Fe-VH interface confirming the good adhesion of the VH phase to Fe particles/matrix after CS (**Fig. 2b**). We also observed regions of interface delamination (**Fig. 2e**), though they were accompanied by cracks in the VH phase. Most likely, VH cracking, discussed above, initiates delamination between Fe and VH phases upon CS. It is worth noting that with the exception of noble metals, metallic surfaces are always passivated by an oxide film at ambient conditions (air, atmospheric pressure, RT) of several nanometers thickness [48]. VH primarily interacts with the Fe oxide film that covers the Fe particles during mixing of the powders and the initial compaction stage (prior to plastic deformation of the metal). Fe-VH interaction takes place only when Fe lattice breaks through the Fe oxide film under the applied pressure. Thus, the Fe-VH interface also contains broken Fe oxide films. As we observe good adhesion at Fe-VH interfaces (**Fig. 2c-e**), we assume good adhesion of VH to both Fe and Fe oxides, when cold-sintered at 2.5 GPa, and RT. However, scanning transmission electron microscopy (STEM) would provide more details on the structure of Fe-VH interface, composition of Fe oxide films and their interaction with VH.

### 3.3. Density and mechanical properties of VH-free and VH-loaded Fe and Fe-Fe_2_O_3_

Orthopaedic implants are exposed to compressive stress. In this framework, we characterized the relative density (expressed as %) and the mechanical properties of the different specimens under compression. Values of these parameters for VH-free and VH-loaded (1 wt%) Fe and Fe-Fe_2_O_3_ are summarized in **Table 1**. Representative HR-SEM micrographs (BSE detector) of VH-loaded Fe and Fe-10Fe_2_O_3_ surfaces are shown in **Fig. 3**. Light-grey and black colors in **Fig. 3a** correspond to Fe and VH phases, respectively. Light-grey, dark-grey and black colors in **Fig. 3b** correspond to Fe, Fe_2_O_3_ and VH phases, respectively.

**Fig. 3.**
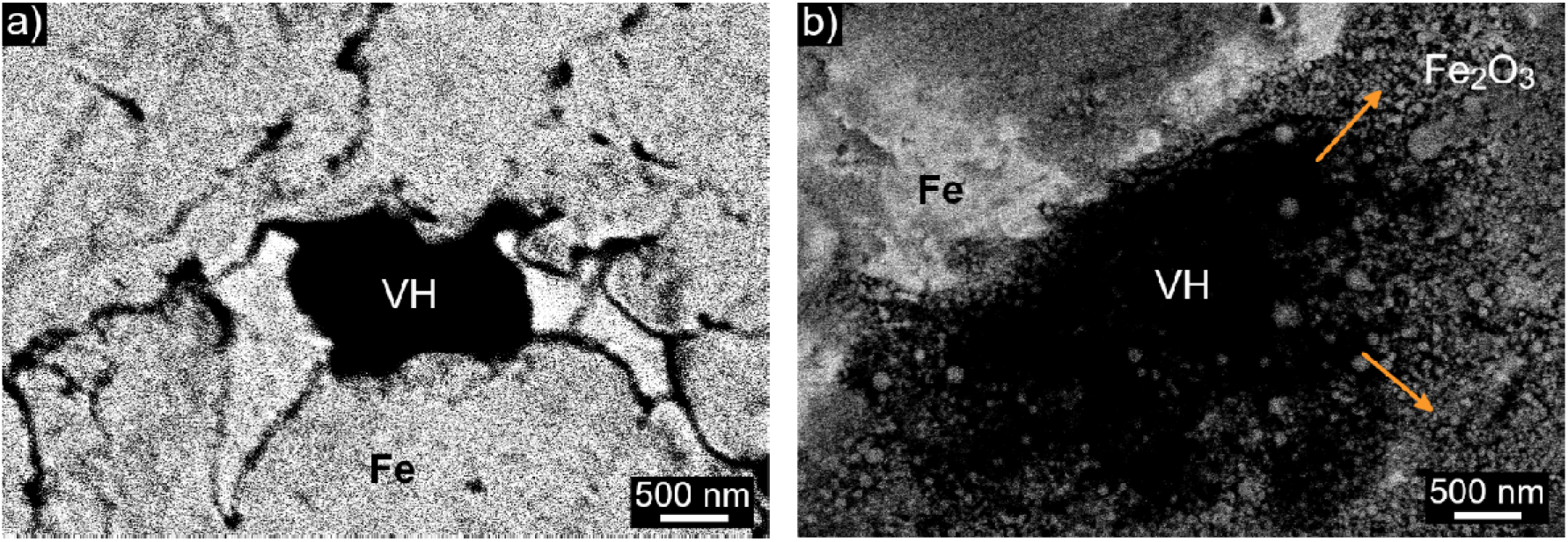
HR-SEM micrographs (BSE detector) of sample surfaces: (a) VH-loaded (1 wt%) Fe and (b) VH-loaded (1 wt%) Fe-10Fe_2_O_3_. Phases of VH, Fe and Fe_2_O_3_ are visualized in black, light-grey and dark-grey colors, respectively. Arrows in (b) point towards non-sintered Fe_2_O_3_ particles. Arrows in (b) point towards Fe_2_O_3_ nanoparticles remained unsintered in Fe-10Fe_2_O_3_ composite cold-sintered at 2.5 GPa, at RT.

**Table 1.** Density, compressive strength, and degradation rate of VH-free and VH-loaded (1 wt%) Fe, Fe-Fe_2_O_3_ produced in this work.

| Material <sup>a</sup> | Relative density,<br>% ± S.D. | Compressive strength,<br>MPa ± S.D. |  | Degradation rate,<br>mm/year ± S.D. |
| --- | --- | --- | --- | --- |
| | | $\sigma_y$ | $\sigma_{ult}$ | |
| Fe | 98.45 ± 0.92 | 540.0 ± 17.9 | 890.0 ± 10.2 | 1.01 ± 0.04 |
| Fe-1wt% VH | 97.37 ± 0.73 | 481.0 ± 10.1 | 584.0 ± 15.1 | - |
| Fe-5Fe<sub>2</sub>O<sub>3</sub> | 94.32 ± 1.77 | 584.0 ± 17.7 | 873.0 ± 53.1 | 1.24 ± 0.08 |
| Fe-5Fe<sub>2</sub>O<sub>3</sub>-1 wt% VH | 96.81 ± 0.47 | 604.0 ± 19.4 | 780.0 ± 13.1 | - |
| Fe-10Fe<sub>2</sub>O<sub>3</sub> | 93.48 ± 2.02 | 600.0 ± 4.0 | 896.0 ± 10.0 | 1.22 ± 0.02 |
| Fe-10Fe<sub>2</sub>O<sub>3</sub>-1 wt% VH | 94.96 ± 2.01 | 604.0 ± 27.4 | 774.0 ± 23.4 | - |
| Fe-15Fe<sub>2</sub>O<sub>3</sub> | 93.07 ± 1.24 | 571.0 ± 44.6 | 819.0 ± 19.8 | 1.18 ± 0.09 |

<sup>a</sup> Drug loading was performed for Fe, Fe-5Fe<sub>2</sub>O<sub>3</sub> and Fe-10Fe<sub>2</sub>O<sub>3</sub> due to their high density and mechanical strength.

Samples produced from microstructured Fe powders show high density and mechanical strength when CS at 2.5 GPa, RT exceeding that one of Fe prepared from nanostructured powders [25]. Relative density of cold-sintered pure Fe from microstructured powder is 98.45 ± 0.92% (**Table 1**), while from pure Fe nanostructured powder is 97.0 ± 0.5% [25]. With addition of Fe_2_O_3_ nanopowders, the relative density of Fe-Fe_2_O_3_ composites decreases. The lowest value of relative density was observed for Fe-15Fe_2_O_3_ composites, 93.07 ± 1.24% (**Table 1**). The decrease is explained by the lack of Fe_2_O_3_ sintering under 2.5 GPa, at RT [49,50], because the processing conditions are insufficient to plastically deform Fe_2_O_3_ particles. Fe_2_O_3_ phase in Fe-Fe_2_O_3_ composites is agglomerations of Fe_2_O_3_ nanoparticles rather than consolidated grains. This is clearly seen at the surface of Fe-10Fe_2_O_3_, where Fe_2_O_3_ nanoparticles are separated from each other (arrows in **Fig. 3b** pointing to dark grey spheres of Fe_2_O_3_ nanoparticles), unlike plastically deformed Fe particles that form sintered matrix (light-grey phase in **Fig 3**). As result, addition of Fe_2_O_3_ lowers matrix integrity, and amount of Fe_2_O_3_ nanopowders negatively correlates with the value of the relative density (**Table 1**).

As expected, addition of 1 wt% VH decreases the relative density of metals and metal-ceramic composites, to 97.37 ± 0.73%, 96.81 ± 0.47%, 94.96 ± 2.01% for Fe, Fe-5Fe_2_O_3_ and Fe-10Fe_2_O_3_, respectively (**Table 1**). Nonetheless, these values are significantly higher than the relative density of Fe10Ag with 1 wt% VH (91.80 ± 0.63%) prepared from the nanostructured Fe10Ag powders using the same processing conditions [24]. This increase confirms our conceptual approach that microstructured metallic powders can possess densification of drug-eluting metals and metal-ceramic composites produced by CS. Surprisingly, VH slightly increases the relative density of drug-eluting Fe-5Fe_2_O_3_ and Fe-10Fe_2_O_3_ (in both cases to ~1.5%, **Table 1**). Most probably a small VH amount improves the integrity of the Fe_2_O_3_ phase due to the good adhesion of VH to Fe oxides, as discussed above. In other words, the drug would play the role of a “glue” that binds Fe_2_O_3_ nanoparticles resulting into small increase of relative density of drug-eluting metal-ceramic composites. Under compression, Fe and Fe-Fe_2_O_3_ show very high strength (**Table 1**) comparable to Fe and 316L stainless steel produced by conventional manufacturing methods [8]. Representative stress-strain curves of VH-free Fe and Fe-Fe_2_O_3_ are shown in **Fig. 4** (grey plots). Pure Fe shows compressive yield (*σ_y_*) and ultimate (*σ_ult_*) strengths of 540.0 ± 17.9 MPa and 890.0 ± 10.2 MPa, respectively (**Table 1**). In general, the addition of a small amount of Fe_2_O_3_ nanoparticles strengthens Fe matrix. For example, 5Fe_2_O_3_ and 10Fe_2_O_3_ increases *σ_y_* to 584.0 ± 17.7 MPa and 600.0 ± 4.0 MPa, respectively, along with the insignificant increase in *σ_ult_* (**Table 1**). While addition of 15Fe_2_O_3_ still shows increase in *σ_y_* to 571.0 ± 44.6 MPa, there is a pronounced decrease in the *σ_ult_* to 819.0 ± 19.8 MPa compared to Fe-5Fe_2_O_3_ and Fe-10Fe_2_O_3_ composites (**Table 1**). The strengthening effect in addition of small amounts of Fe_2_O_3_ (5 and 10 vol%) is explained by the brittle nature of Fe_2_O_3_ nanoparticles that constrain the plastic flow of Fe matrix, which increases the *σ_y_* of Fe under the compressive load. Further increase of the Fe_2_O_3_ amount to 15 vol.% decreases the mechanical strength of Fe matrix due to the low integrity of Fe_2_O_3_ phase discussed previously (**Table 1**). Representative stress-strain curves show increase of Fe matrix *σy* and stiffness in addition of 5Fe_2_O_3_ and 10Fe_2_O_3_ (grey curves in **Fig. 4**, steeper slopes of elastic region), which reflects the strengthening effect by Fe_2_O_3_ nanoparticles. However, addition of brittle Fe_2_O_3_ nanoparticles decreases plasticity of Fe matrix, which can be seen in sharper decline of plastic regions with increase of Fe_2_O_3_ (grey curves in **Fig. 4**). Based on these results, we selected compositions with higher density and mechanical strength for the VH loading, i.e., Fe, Fe-5Fe_2_O_3_ and Fe-10Fe_2_O_3_.

**Fig. 4.**
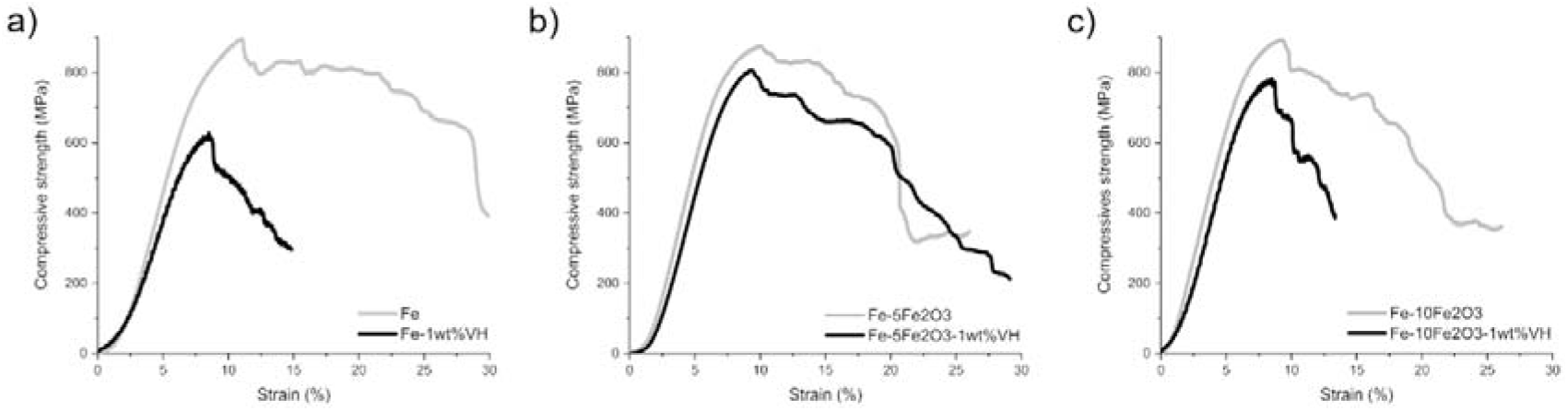
Representative stress-strain curves of VH-free and VH-loaded (1 wt%) samples under compression: (a) Fe, (b) Fe-5Fe_2_O_3_ and (c) Fe-10Fe_2_O_3_. Grey and black curves correspond to VH-free and VH-loaded samples, respectively.

Representative stress-strain curves of VH-loaded (1 wt%) Fe and Fe-Fe_2_O_3_ are shown in **Fig. 4** (black plots). As expected, addition of VH decreased the mechanical performance of the materials (**Fig. 4**). The most pronounced decrease upon VH loading was observed for pure Fe with a *σ_y_* drop from 540.0 ± 17.9 to 481.0 ± 10.1 MPa (**Table 1**). Interestingly, addition of VH slightly increased *σ_y_* of Fe-5Fe_2_O_3_ and Fe-10Fe_2_O_3_ composites, both to approximately 604.0 MPa (**Table 1**). We attribute this behavior to the enhanced stability of Fe_2_O_3_ agglomerates due to the “glue effect” of VH, as previously discussed. However, *σ_ult_* in VH-loaded Fe, Fe-5Fe_2_O_3_ and Fe-10Fe_2_O_3_ decreased to 584.0 ± 15.1 MPa, 780.0 ± 13.1 MPa and 774.0 ± 23.4 MPa, respectively (**Table 1**). Decrease of *σ_ult_* upon incorporation of VH is more pronounced for pure Fe than for Fe-Fe_2_O_3_ composites (**Fig. 4**). It is most likely because of the strengthening effect stemming from Fe_2_O_3_ agglomerates (grey plots in **Fig. 4**), which stability is additionally enhanced by the small amount of VH. The mechanical strength of VH-loaded Fe-Fe_2_O_3_ composites was approximately 2.5-times higher than of previously reported VH-loaded Fe10Ag nanocomposites [24]. Although these materials are different in the secondary phase (Fe_2_O_3_ versus Ag), the volume of the primary Fe phase is comparable (90 ± 5 vol%). Moreover, the following process parameters are identical: chemical composition of starting Fe powder (carbonyl Fe), drug composition and amount (1 wt% VH), loading procedure (manual mixing) and consolidation conditions (2.5 GPa, at RT). The difference between VH-loaded Fe10Ag and Fe-Fe_2_O_3_ composites is in the microstructure of Fe powders, which stems from different preparation routes of Fe10Ag and Fe-Fe_2_O_3_ powders prior to the drug-loading. In the case of Fe10Ag, microstructured carbonyl Fe and Ag_2_O nanopowders undergo high-energy attrition milling, which refines them to nanostructured particles [24]. In the case of Fe-Fe_2_O_3_, microstructured carbonyl Fe was manually blended with Fe_2_O_3_ nanopowders. Such procedure does not imply structural changes and preserves the original microstructure of carbonyl Fe. Compressive *σ_y_* of VH-loaded Fe10Ag is 180 ± 12 MPa [24], while *σ_y_* of VH-loaded Fe and Fe-10Fe_2_O_3_ prepared in this work is 481.0 ± 10.1 and 604.0 ± 27.4MPa, respectively. These results confirm that the microstructure of the primary metallic phase plays the key role in mechanical strength of drug-eluting metals and metal-ceramic composites produced by CS (2.5 GPa, at RT).

High mechanical strength of drug-eluting Fe and Fe-Fe_2_O_3_ makes them promising candidates for load-bearing bone healing. The mechanical strength of the materials produced in this work is ~3 times higher than the maximum strength of the cortical bone (~220 MPa) [29,30]. Superior mechanical performance would enable decreasing the amount of material required for temporary support at the healing site. This can be realized through the use of smaller/thinner implants, or high-porous scaffolds [51]. Moreover, superior mechanical strength and low processing temperature would enable further modifications of the matrix composition to support bone regeneration, e.g., by adding thermosensitive bioactive molecules (i.e., BMPs), biopolymers, or bio-glasses. Furthermore, the simplicity of the proposed processing route expands the opportunities for the design of LDD therapies at load-bearing body sites. Since the drug-loading mechanism is based on a physical phenomenon (plastic deformation), it does not imply chemical reaction and ensures near 100% encapsulation efficiency of one or more drugs as opposed to other methods such as infiltration [52] or impregnation [53]. Finally, stronger metals or metal-ceramic composites would enable the loading of more drug without jeopardizing them. Synergy effect of added materials, bioactive molecules, and loaded drugs might significantly support bone restoration at the healing sites [54].

### 3.4. Degradation of Fe and Fe-Fe_2_O_3_

While addition of Fe_2_O_3_ increased mechanical performance of Fe matrix (**Table 1**), the main goal of Fe_2_O_3_ incorporation was to increase the degradation rate of Fe. Fe_2_O_3_ acts as cathode, enhancing dissolution of anodic Fe, as previously reported for spark-plasma-sintered Fe-Fe_2_O_3_ composites [39] and X-70 steels [55]. To confirm that for Fe-Fe_2_O_3_ composites prepared by CS, we assessed the relative weight loss of VH-free samples during eight weeks of in MH solution (**Fig. 5**). pH of the MH solution slightly increased to pH 7.5 for all the samples during the assay.

**Fig. 5.**
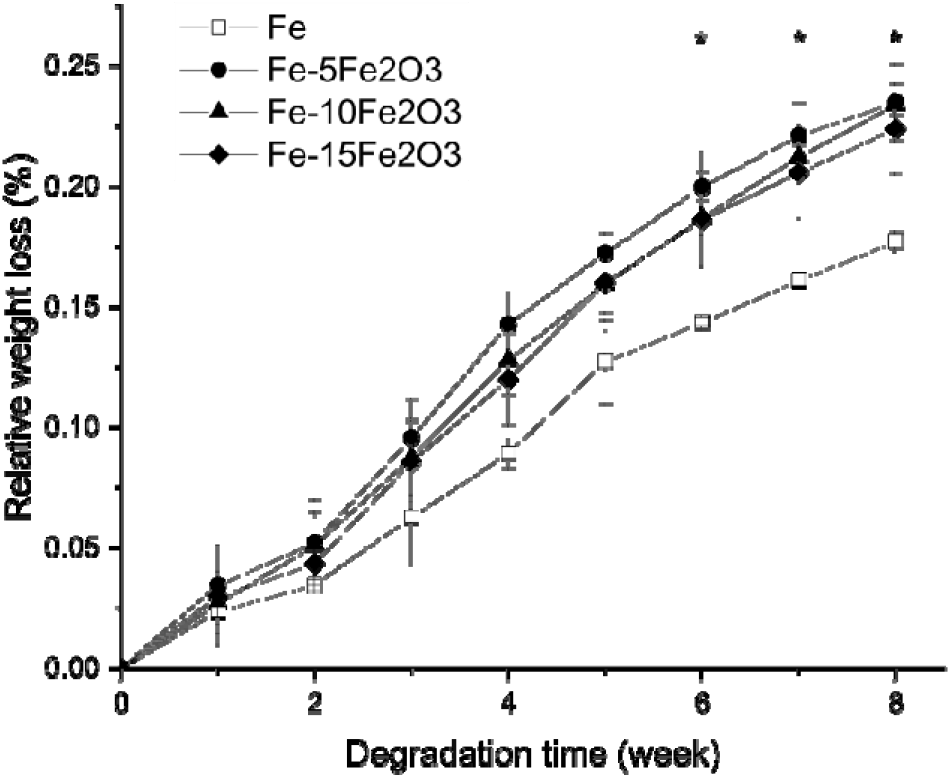
Relative weight loss of Fe and Fe-Fe_2_O_3_ composites as a function of the degradation (immersion) time. * Statistically significant difference (p < 0.05) between relative weight losses of Fe and each of Fe-Fe_2_O_3_ composites starts from week 6.

Addition of Fe_2_O_3_ increased the degradation rate of Fe matrix from 1.01 ± 0.04 mm/year to 1.24 ± 0.08 mm/year and 1.22 ± 0.02 mm/year for Fe-5Fe_2_O_3_ and Fe-10Fe_2_O_3_ composites, respectively (**Table 1**). Addition of 15Fe_2_O_3_ had minor increase of the degradation rate to 1.18 ± 0.09 mm/year (**Table 1**). Significant difference between relative weight losses of Fe and each of Fe-Fe_2_O_3_ composites was observed only after week 6 (**Fig. 5**). Increase in the degradation rate by the small amount of Fe_2_O_3_ (5 vol%) and following decline adding higher Fe_2_O_3_ amounts (10, 15 vol%) is in good agreement with previously reported results on the degradation rate of spark-plasma-sintered Fe-Fe_2_O_3_ [39]. This trend might be explained by the semi-conductive nature of Fe_2_O_3_, which obstructs the electron transfer within the Fe matrix required for the micro-galvanic corrosion. Moreover, a decline in the corrosion rate with higher amount of Fe_2_O_3_ might relate to the corrosion layer formed at the initial stage of degradation. High amount of cathodic Fe_2_O_3_ (10, 15 vol%) might accelerate formation of corrosion layer right after the immersion, which hampers following access of anodic Fe to the degradation medium.

### 3.5. Antibacterial activity of VH-free and VH-loaded Fe and Fe-10Fe_2_O_3_

To confirm the release of VH from the developed drug-eluting Fe and Fe-10Fe_2_O_3_ composites at antibacterial concentrations, we performed ZOI tests against *S. aureus* (**Fig. 6**). For this, we collected extracts from the VH-loaded Fe and Fe-10Fe_2_O_3_ over 14 days. VH-free samples were used as controls.

**Fig. 6.**
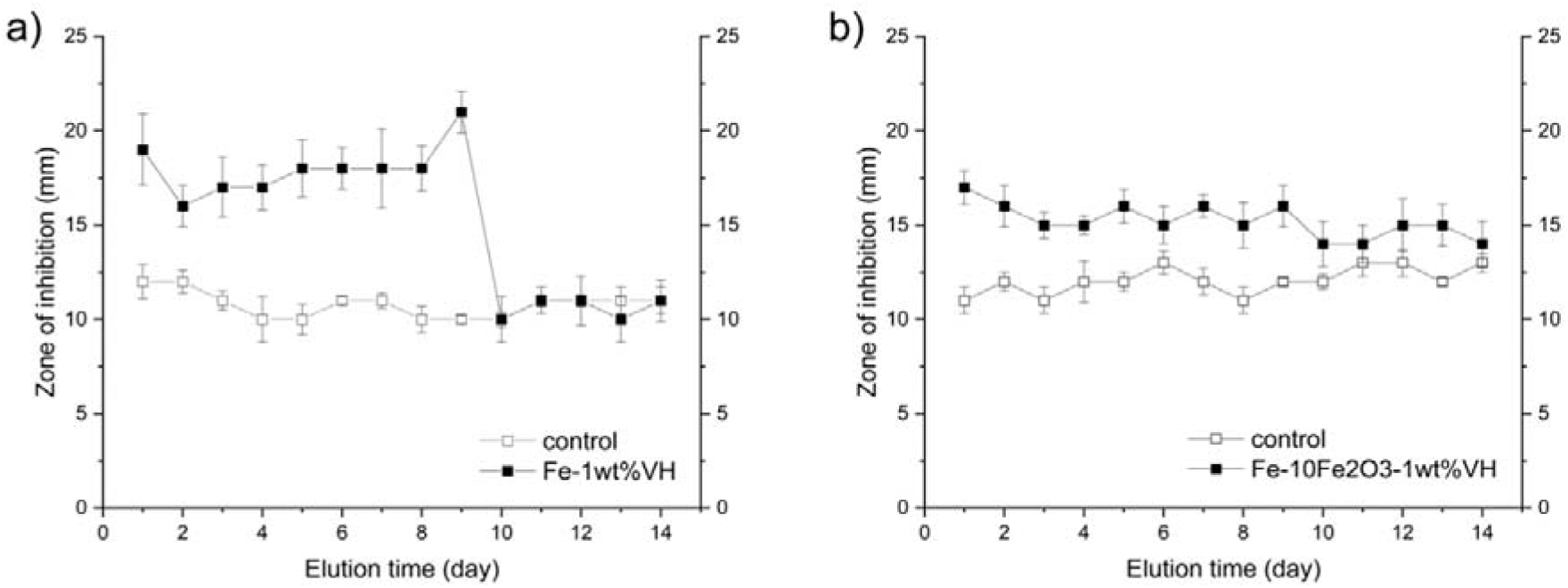
ZOI test against *S. aureus* of VH-loaded (1 wt%): (a) Fe and (b) Fe-10Fe_2_O_3_.

Pure Fe and Fe-10Fe_2_O_3_ showed antibacterial activity against *S. aureus* with ZOI of 11.0 ± 0.3 and 12.0 ± 0.2 mm, respectively (**Fig. 6**), which exceeds the diameter of test disk (6 mm). Antibacterial properties of the controls most probably result from the degradation of the Fe. Series of Fe oxidation-reduction reactions cause formation of different reactive oxygen species (ROS) such as OH and HO_2_ [56]. ROS-induced oxidative stress leads to damage of the bacterial membrane and bacterial death. This also explains the larger ZOI obtained for Fe-10Fe_2_O_3_ controls compared to Fe controls (**Fig. 6**); Fe_2_O_3_ increases Fe degradation and the ROS concentration.

Loading of VH (1 wt%) into Fe samples led to an increase of the ZOI for 9 days (**Fig. 6a**), which indicates drug release during this period. At day 1, we observed burst release corresponding to VH dissolution from sample surface due to its high aqueous solubility in broad pH range. During days 2-8, the ZOI slightly increased with a more pronounced increase in day 9 (**Fig. 6a**). There are two factors that might contribute to the ZOI increase: (i) more efficient VH release and (ii) release of ROS generated from the Fe matrix degradation. After the burst release (day 1), the drug dissolves in the open pores and channels. A further increase of the ZOI could be ascribed to the higher amount of drug stored in the inner pores compared to those at the surface. Moreover, drug dissolution and release generate new pores and increases the surface area to a greater extent than in denser controls (**Table 1**), which can also result in the generation of greater ROS concentrations. After day 10, the ZOI of VH-free and VH-loaded Fe samples was similar (**Fig. 6a**), meaning that no drug was released after day 9. Initial release of antibiotic drug can be a promising path to prevent bacterial contamination on bulk metallic and metal-ceramic implant and/or in surrounding tissues after implantation. In the case of Fe-10Fe_2_O_3_ samples loaded with the same amount of VH (1 wt%), we observed a different ZOI trend (**Fig. 6b**). Unlike VH-loaded Fe, the drug release from Fe-10Fe_2_O_3_ was prolonged for up to 14 days, as confirmed by the antibacterial activity of the release medium (**Fig. 6b**). In general, ZOI of drug-eluting Fe-10Fe_2_O_3_ was lower and gradually decreased during the test period. The amount of drug and loading procedure were identical for both materials. Thus, these results strongly suggest that the addition of 10Fe_2_O_3_ to the Fe matrix slows down the initial drug dissolution. This result looks counterintuitive, as increased degradation rate of Fe-10Fe_2_O_3_ (**Table 1**) should promote drug release from open pores. However, weight losses of Fe and Fe-10Fe_2_O_3_ during the first two weeks has statistically insignificant difference (**Fig. 5**). At the same time, faster degradation rate can promote formation of corrosion products after immersion, which clog pores and obstruct drug outer diffusion to the release medium. Therefore, increased degradation rate slows down/extends drug release at the initial stage. For better understanding of how matrix degradation rate influences on drug release, significant degradation of the matrix is required. According to our degradation studies (**Fig. 5**), this investigation should take longer than six weeks, when significant difference between degradation rates of Fe and Fe-10Fe_2_O_3_ is observed.

### 3.6. Microstructure of VH-loaded Fe after VH release

We suggested that drug release from bulk metals and metal-ceramic composites is a two-stage process: (i) initial release from surface and adjacent open pores, and (ii) delayed release of drug from the matrix upon its degradation. The low antibacterial activity observed in the ZOI test indicates the end of the initial release stage (**Fig. 6**). To very that, we performed FIB analysis of surface and the bulk of VH-loaded Fe after two weeks of immersion/drug release. We performed immersion test of VH-loaded Fe in conditions identical to degradation tests (see Section 3.4) for the period identical to ZOI tests (see Section 3.5). To access the sample bulk, we polished it on 1200-grit silicon carbide paper. The thickness removed by polishing (investigated depth) was ~240 μm.

HR-SEM micrographs of VH-loaded Fe after two weeks of immersion/drug release in MH solution are summarized in **Fig. 7**. The sample after polishing is shown in **Fig. 7a**.

**Fig. 7.**
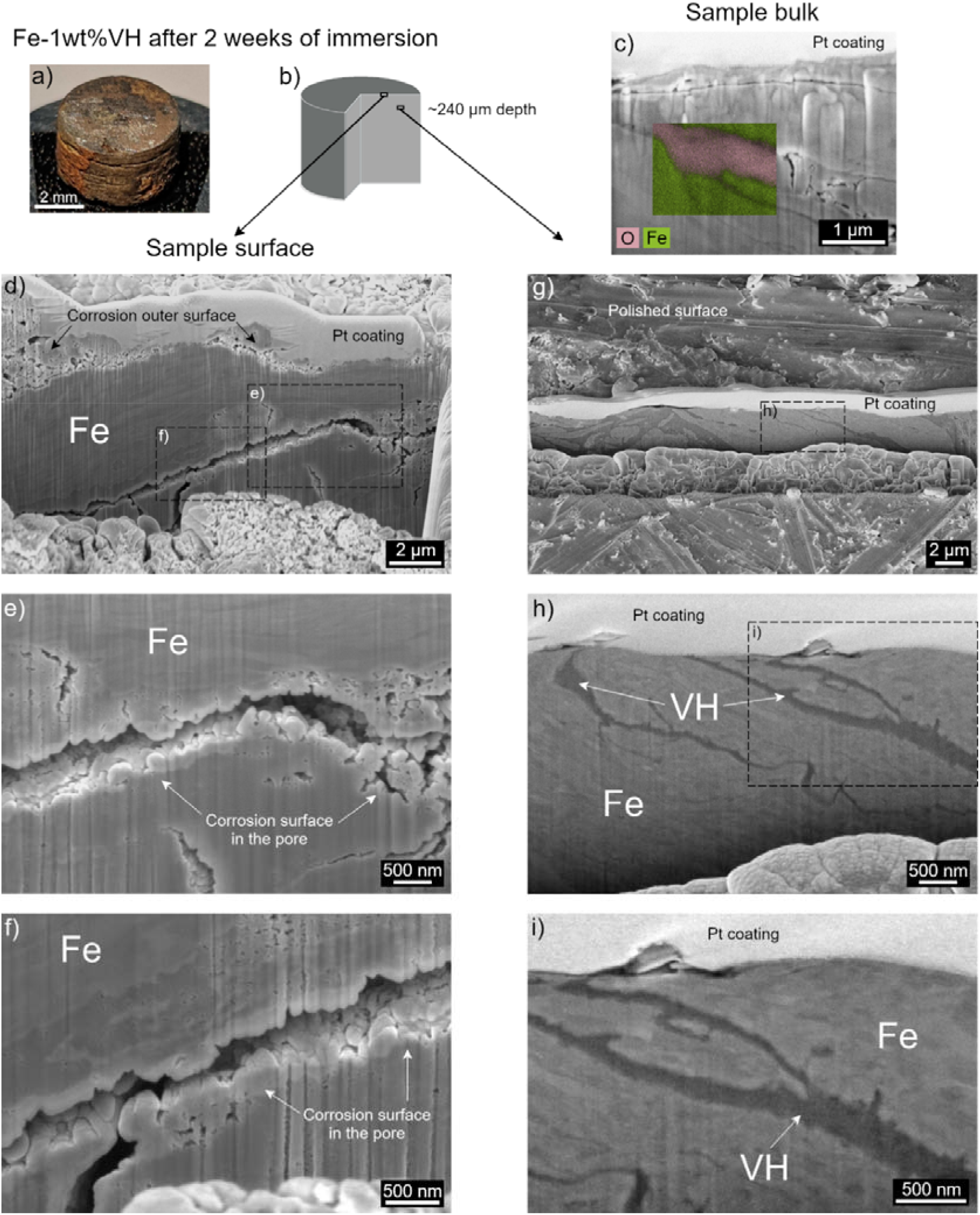
VH-loaded (1 wt%) Fe after two weeks of immersion/drug release in MH solution: (a) sample photo; (b) illustrative isometric projection of the sample showing FIB-investigated regions; (c) HR-SEM micrograph with elemental map of the inner sample (~240 μm depth); (d-f) HR-SEM micrographs of sample cross-section at the surface, (g-i) HR-SEM micrographs of the inner sample at ~240 μm depth. Pt coating – protective coating for FIB milling. Vertical texturing of the cross-sectional surfaces (c, e, f) corresponds to FIB “curtaining effect”.

Isometric projection of the sample in **Fig. 7b** illustrates the investigated areas. **Fig. 7d-f** show HR-SEM micrographs of surface cross-sections and **Fig. 7g-i** provide HR-SEM micrographs of inner cross-sections at different magnifications. Elemental map of the sample bulk after two weeks of immersion is provided in **Fig. 7c**. After two weeks of immersion, we observed no drug at the surface and in the adjacent pores (**Fig. 7d-f**). Roughness and porosity at the surface correspond to corrosion (**Fig. 7d**). Adjacent pores had analogous corrosion surface (**Fig. 7d-f**). Texturing of the cross-section (better seen in **Fig. 7e,f**) corresponds to the “curtaining effect” from FIB milling (see Supplementary Materials, **Fig. S2**). Elongated shape of the pore is similar to the shape of VH phase loaded into Fe matrix (**Fig. 2**), which indicates that this pore contained VH prior its dissolution during the initial stage of the release. HR-SEM of the pore show increased surface roughness and corrosion products inside the pores (**Fig. 7e**). Since corrosion products obstruct the access of the medium to the pore, they can hamper VH diffusion at least during the initial stage of the release. Consequently, an increase of matrix degradation rate will increase corrosion buildups in pores and slow down/extend the drug release. We observed this effect in ZOI tests, when increased degradation rate of Fe matrix by adding 10Fe_2_O_3_ slowed down drug release and extended it from 9 to 14 days (**Fig. 6**).

HR-SEM micrographs of the inner regions of VH-loaded Fe after two weeks of immersion are presented in **Fig. 7c,g-i**. HR-SEM and EDS analyses confirm that the drug is present at least at ~240 μm depth even after two weeks of immersion. EDS analysis confirms that grey and black phases in the cross-sectional images correspond to Fe matrix and embedded drug (oxygen-rich phase), respectively (**Fig. 7c**). In addition, the microstructure of VH in the Fe bulk after two weeks of immersion (**Fig. 7g-i**) is similar to the one in the as-prepared sample (**Fig. 2**). These results indicate the termination of the drug release after 1-2 weeks (**Fig. 6**) and confirm that only the drug at the surface and adjacent pores is released (**Fig. 7d-f**). Delayed release of the drug from the inner regions of the material will occur upon metallic matrix degradation, so drug release will positively correlate with the degradation rate of the matrix. However, in the case of Fe and Fe-Fe_2_O_3_ significant difference in the delayed drug release might occur only after six weeks, when the difference between degradation rates of the materials becomes significant (**Fig. 5**).

### 3.7. Cell compatibility of Fe and Fe-10Fe_2_O_3_

To verify that the metallic and metal-ceramic matrices produced in this work exhibit no signs of cytotoxicity, we performed flow cytometry studies of the cold-sintered Fe and Fe-10Fe_2_O_3_ on sensitive fibroblasts (NIH/3T3 line). Results of Fe and Fe-10Fe_2_O_3_ flow cytometry studies are shown in **Fig. 8**. Polystyrene tissue culture plastic was used as control to ensure optimal cell adhesion and proliferation.

**Fig. 8.**
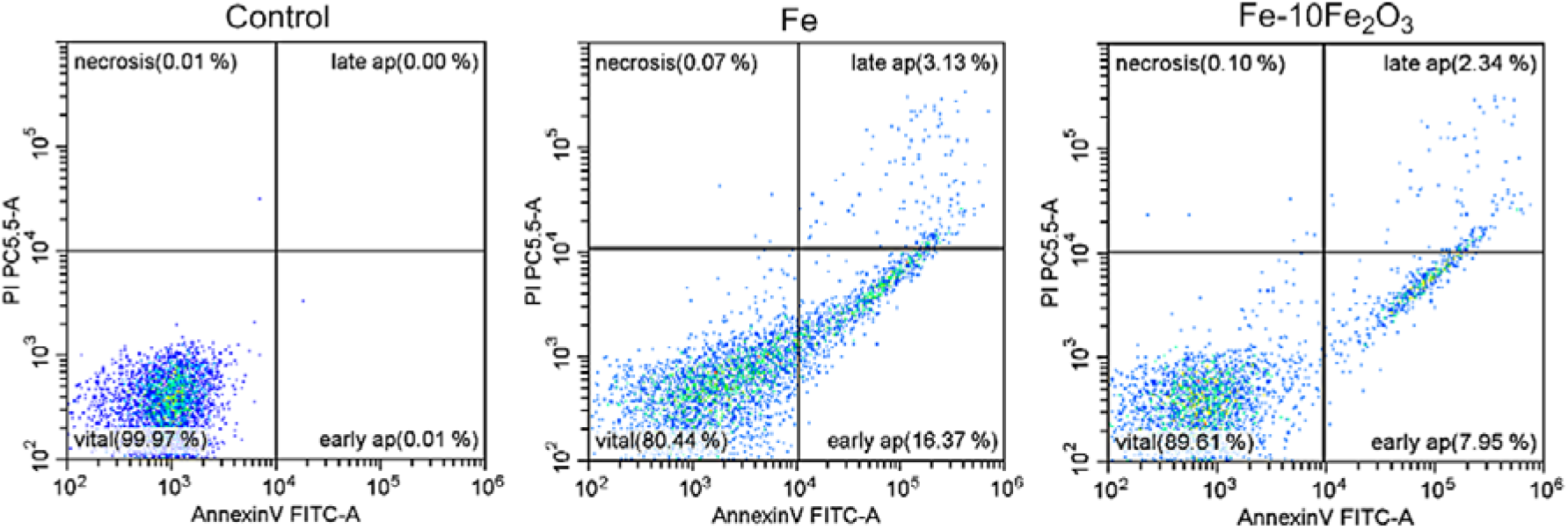
Analysis of the different steps of apoptosis and the necrotic cell percentage of fibroblast NIH/3T3 cultured on tissue culture plastic (Control), on Fe and Fe-10Fe_2_O_3_ composite by flow cytometry. Identification of viable (vital), early apoptotic (early ap), late apoptotic (late ap) and necrotic cells (necrosis) was carried out by propidium iodide staining and forward angle light scatter analysis.

The percentage of apoptotic cells increased from 0.01 % for the control to 7.17 % and 6.36 % for Fe and Fe-10Fe_2_O_3_, respectively (**Fig. 8**). At the same time, no significant difference in the percentage of necrotic cells between the control and Fe, Fe-10Fe_2_O_3_ was observed, which were 0.04 %, 0.04 % and 0.06 %, respectively. The percentage of viable cells on the samples was >90%, which indicated adhesion and proliferation of NIH/3T3 fibroblasts on the surface of microstructured Fe and Fe-10Fe_2_O_3_ prepared by CS and confirmed the good cell compatibility of these materials. These results indicate that microstructured Fe and Fe-10Fe_2_O_3_ produced by CS induce a small increase of apoptosis in NIH/3T3 cell line without causing necrosis.

## 4. Conclusions

LDD via degradation of temporary metallic implants is an attractive concept in bone restoration. In this work we report on characterization of bulk VH-free and VH-loaded Fe and Fe-Fe_2_O_3_ composites produced by CS method at 2.5 GPa, at RT. High mechanical properties of the VH-free materials were achieved using microstructured metallic powder (Fe, 90 ±5 vol%) and small fraction of ceramic nanopowder (Fe_2_O_3_, 10 ±5 vol%). Microstructured metallic powder enables better plastic flow, while ceramic nanopowder strengthens the metallic matrix. Moreover, small amount of Fe_2_O_3_ increases degradation rate of Fe. Cell viability tests on NIH/3T3 fibroblasts shows no toxicity of the microstructured Fe and Fe-10Fe_2_O_3_ composites. High performance of VH-free metals and metal-ceramic composites enabled to obtain very strong VH-loaded (1 wt%) Fe and Fe-Fe_2_O_3_ composites with compressive ultimate strength that is up to 3-fold greater than that of cortical bone. VH-loaded (1 wt%) Fe and Fe-10Fe_2_O_3_ suppressed *S. aureus* growth during first 9 and 14 days, respectively. This antibacterial performance corresponds to initial stage of drug release when drug dissolves from surface and adjacent pores (in scale of 1-2 weeks). Increase in Fe degradation rate slows drug release during the initial stage. One example of VH-loaded Fe we show that drug remains in the bulk of the metal after the initial release. Overall results indicate that the cold-sintered drug-eluting metals and metal-ceramic composites are promising prototypes for the development of temporary implants with LDD features at load-bearing sites because the production method relies on the plastic deformation of metallic particles, and it can be used for the loading of small-molecule drugs, active macromolecules (e.g., BMPs) into other biodegradable metals (magnesium, zinc, molybdenum) or metal-based composites.

## Credit author statement

Conceptualization – A. Sharipova, E. Gutmanas, A. Sosnik, M. Lerner; Funding acquisition – A. Sosnik, E. Gutmanas, M. Lerner; Investigation - A. Sharipova, O. Bakina; Methodology - A. Sosnik, A. Sharipova, E. Gutmanas, O. Bakina, A. Lozhkomoev; Data analysis - A. Sharipova, O. Bakina, A. Sosnik, A. Lozhkomoev; Writing - original draft, review & editing – A. Sharipova, O. Bakina, A. Sosnik. All authors have read and agreed to the submitted version of the manuscript.

## Acknowledgments

This work was financed by the Ministry of Science & Technology, Israel (grant 3-16574) and Russian Foundation for Basic Research (grant 19-53-06006). The partial support of the Russell Berrie Nanotechnology Institute (RBNI, Technion-Israel Institute of Technology) is also acknowledged. A. Sosnik and A. Sharipova thank Dr. K. Zohar-Hauber, E. Naimark, R. Biletsky, E. Strokin and A. Garkun from the Advanced Materials Characterization Lab of the Israel Institute of Metals for their support to make this work possible. A. Sharipova thanks the Minerva Fellowship Program (German Federal Ministry for Education and Research) for financial support. A. Sosnik thanks the support of the Tamara and Harry Handelsman Academic Chair. The partial financial support of the Russell Berrie Nanotechnology Institute (RBNI, Technion-Israel Institute of Technology) is also acknowledged.

