## Supplemental information for "Drug-eluting biodegradable metals and metal-ceramic composites: High strength and delayed drug release"

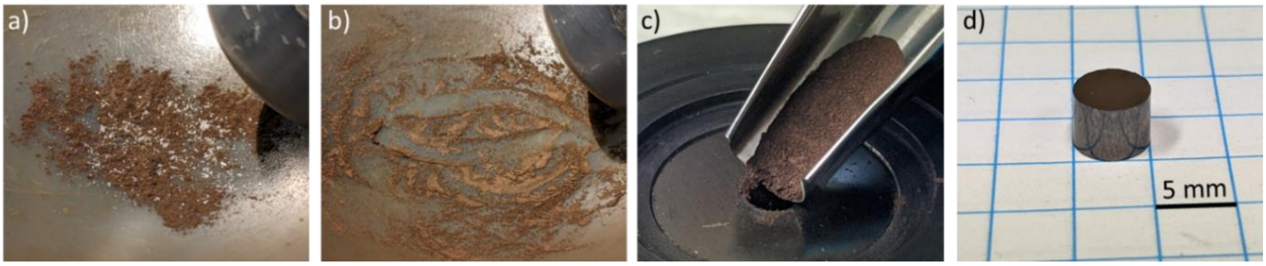


**Fig. S1** – Fabrication of Fe-5$Fe_2O_3$-1wt%VH: (a) Fe-5$Fe_2O_3$ and VH powders are mixed in a mortar before mixing (ginger color and white color correspond to Fe-5$Fe_2O_3$ blend and VH drug, respectively); (b) Fe-5$Fe_2O_3$ and VH powders after dry mixing in mortar; (c) the Fe-5$Fe_2O_3$-1wt%VH blend is loaded to Ø 5 mm consolidation die; and (d) Fe-5$Fe_2O_3$-1wt%VH sample is released from the die.

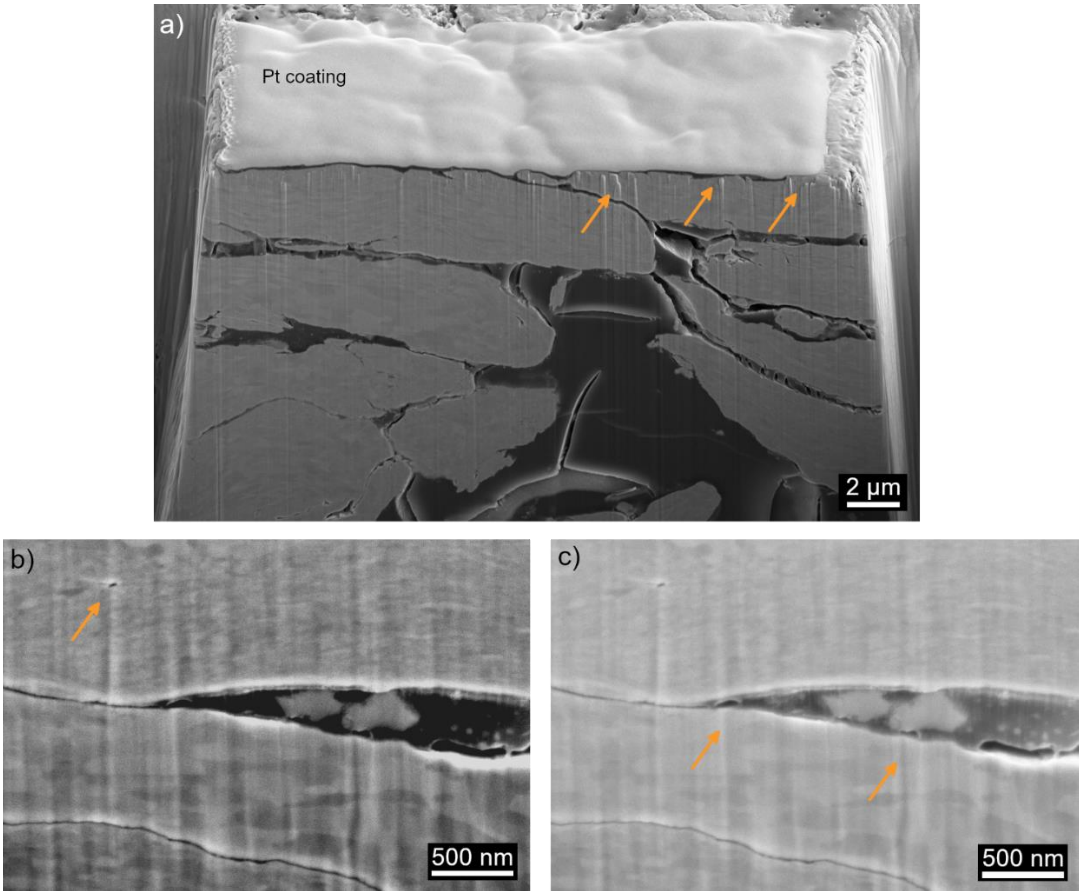


**Fig. S2** - Representative cross-sectional HR-SEM (TLD detector) micrographs of Fe-5wt%VH. Dark and grey colors correspond to VH and Fe phases, respectively. (a) Representative cross-sectional micrograph used for 3D-reconstruction; (b,c) cross-sectional micrograph of Fig. 3c of the manuscript with adjusted brightness-contrast level. Increased contrast in (b) improves the visualization of Fe grain orientation. Increased brightness in (c) improves the visualization of VH microstructure and the Fe-VH interface. The vertical texturing artifacts in the Fe matrix stems from FIB milling and it is called “curtaining effect”. The effect occurs between phases with different milling behavior. In the case of metal loaded with drug, curtaining effect occurs at interfaces: air-metal (a, yellow arrows), pore-metal (b, yellow arrow), as well as drug-metal or drug-metal-pore interface (c, yellow arrows). The ‘curtaining effect’ was considered in 3D-micrograph reconstruction (Vortex module of Avizo 9.5 Software, option to remove curtaining artifacts).
